# Novae: a graph-based foundation model for spatial transcriptomics data

**DOI:** 10.1101/2024.09.09.612009

**Authors:** Quentin Blampey, Hakim Benkirane, Nadège Bercovici, Fabrice André, Paul-Henry Cournède

## Abstract

Spatial transcriptomics is advancing molecular biology by providing high-resolution insights into gene expression within the spatial context of tissues. This context is essential for identifying spatial domains, enabling the understanding of micro-environment organizations and their implications for tissue function and disease progression. To improve current model limitations on multiple slides, we have designed Novae (https://github.com/MICS-Lab/novae), a graph-based foundation model that extracts representations of cells within their spatial contexts. Our model was trained on a large dataset of nearly 30 million cells across 18 tissues, allowing Novae to perform zero-shot domain inference across multiple gene panels, tissues, and technologies. Unlike other models, it also natively corrects batch effects and constructs a nested hierarchy of spatial domains. Furthermore, Novae supports various downstream tasks, including spatially variable gene or pathway analysis and spatial domain trajectory analysis. Overall, Novae provides a robust and versatile tool for advancing spatial transcriptomics and its applications in biomedical research.

## 1 Introduction

Spatial transcriptomics^1,2^ data provide invaluable insights into cellular interactions within their micro-environment and the complexities of tissue organization. A key advantage over current single-cell RNA sequencing (scRNAseq)^3^ is that spatial transcriptomics maintains the spatial context of cells, enabling a deeper understanding of how cells interact within their native environments. Technologies in spatial transcriptomics can be broadly categorized into two types: (i) Next-Generation Sequencing^4^ (NGS)-based methods that offer whole-genome sequencing, and (ii) imaging-based techniques like the Xenium^5^, MERSCOPE^6^, or CosMX^7^, that provide subcellular resolution. The former allows for comprehensive gene analysis but lacks fine spatial detail, while the latter offers detailed spatial resolution but with a limited gene panel size. As imaging-based technologies continue to evolve, they expand their gene panel capabilities, enabling the inclusion of larger panels or the replacement of low-quality genes during studies. However, this flexibility often results in experiments conducted across different machines or using varying panels, which introduces new challenges. In more general cases, when performing analysis over multiple spatial transcriptomics slides (for both NGS or imaging-based techniques), it is common to observe a strong batch effect, such that it can be challenging to identify common spatial patterns across multiple slides without careful consideration of the batch effect.

A key focus in spatial transcriptomics is the identification and categorization of spatial micro-environments, often referred to as spatial domains or niches. Various methods, such as STAGATE^8^, GraphST^9^, SpaceFlow^10^, and SEDR^11^, have been developed for this purpose. While these methods show promising results (especially with NGS-technologies with spot resolution like Visium), they are limited by (i) their reliance on predefined gene panels, (ii) sensitivity to batch effects, and (iii) dependence on external tools like Harmony^12^ for batch effect correction and Leiden^13^ or Mclust^14^ for clustering. These dependencies can slow down processing and reduce robustness, since external tools need to be re-run for each new analysis or when adjusting spatial domain resolutions (i.e., choosing different number of spatial domains). Additionally, due to their reliance on specific gene sets, these methods often necessitate training on the intersection of gene sets, which can significantly reduce the number of available genes and, consequently, impact performance. Most importantly, even when applied to slides with a shared panel, these models tend to identify primarily slide-specific domains, which limits the comparison of domains across a broader study and reduces the potential for discovering new spatial biomarkers.

To address these limitations, we introduce Novae, a self-supervised^15^ graph attention network^16^ that encodes local environments into spatial representations. Unlike existing methods, the same Novae model can operate with multiple gene panels, allowing for the application across diverse technologies and tissues. It includes native batch effect correction methods, directly correcting for variations and enhancing robustness and scalability. Therefore, Novae’s design allows it to seamlessly integrate data from different platforms and gene panels without compromising performance. We trained Novae on a large dataset comprising 78 slides, representing nearly 30 million cells across 18 tissues and three different subcellular resolution technologies (Xenium^5^, MERSCOPE^6^, CosMX^7^). This broad training allows Novae to compute relevant representations via zero-shot^17^ or fine-tuning on any new slide from any tissue. These representations can be directly used for spatial domain identification, eliminating the need for external clustering tools. Beyond spatial domain identification, these representations can be applied to various downstream tasks, including (i) spatial domain trajectory analysis, (ii) spatially variable gene analysis, and (iii) spatial pathways analysis.

Novae’s versatility, robustness and ease of use make it a powerful tool for advancing spatial transcriptomics research within the scientific community.

## 2 Results

### 2.1 A foundation model for spatial transcriptomics

Novae is a graph-based model that we trained on a large single-cell spatial transcriptomics dataset composed of about 30 millions cells. This model, aware of 18 different tissues, is hosted on a public hub and can be reused by the user for multiple downstream tasks on any technology or gene panel, as illustrated in Figure 1a. The main application of Novae is to learn spatial domains, which can be performed with the shared model without any re-training (called zero-shot inference^17^). If desired, the user can also re-train Novae to obtain refined results (fine-tuning). Two distinguishing properties of Novae are that (i) it provides a nested organization of spatial domains for different resolutions and (ii) it natively corrects batch effect across slides. This consistent domain assignment across slides enables comparison analyses of a study containing multiple slides. Additionally, Novae can perform multiple downstream analysis tasks, such as (i) spatial pathway analysis, (ii) spatial domain organization/architecture analysis, or (iii) spatial variable genes analysis. These tasks and properties are summarized in Figure 1b.

**Figure 1:**
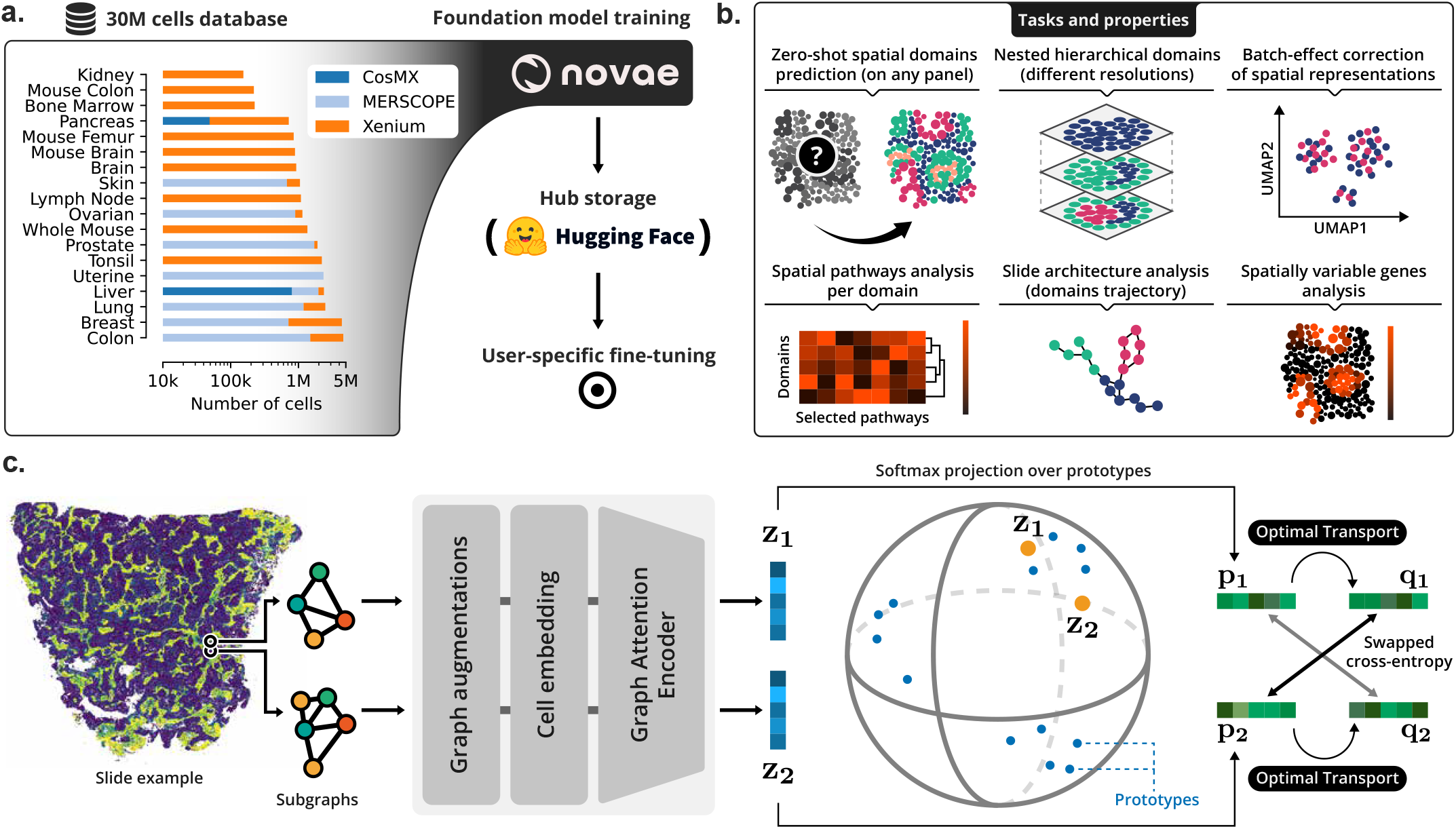
Overview of Novae. **a**. The dataset is composed of 29 million cells across 18 different tissues for three different single-cell spatial transcriptomics technologies. This dataset was used to train Novae, which was shared online on Hugging Face Hub. Users can easily load the pre-trained model for reuse or fine-tuning. **b**. A visual summary of the key tasks and capabilities of Novae. Notably, Novae can infer spatial domains in a zero-shot manner, organizing them hierarchically. Additionally, based on learned representations, Novae can align representations across different batches. Other downstream tasks include spatial patway analysis, slide architecture analysis, and spatially variable gene analysis. **c**. An overview of Novae’s underlying method. The process involves extracting two nearby subgraphs of cells (or spots), which are then augmented and embedded. These graph inputs are processed through a graph attention encoder, generating representations for both subgraphs. These representations are projected onto prototypes—learnable vectors from the latent space. Finally, an optimal transport task is applied to the assignment probabilities over these prototypes, and a swapped cross-entropy loss is computed for backpropagation.

In terms of technical details, Novae is a graph-based neural network that is trained in a self-supervised manner based on the SwAV^18^ framework. It learns representations of the local microenvironment at the single-cell (or spot) resolution. More specifically, for each cell or spot, a representation of its neighborhood is provided by a Graph Attention Network^16^, after embedding each cell into a panel-invariant representation. In order to effectively assign domains, we learn embeddings in the latent space, called prototypes, which represent elementary spatial domains. Depending on the desired level of resolution, these elementary domains are regrouped to form the domains at the desired resolution. We project the spatial-domain representations into these prototypes to get probabilities of assignments. These probabilities are corrected via an algorithm of Optimal Transport^19^ to ensure a smooth representation of the different prototypes. Afterwards, we use cross-entropy loss to make the representation of two cells in the same domain closer together. This methodological approach is illustrated in Figure 1c.

### 2.2 Pan-tissue spatial domains

As outlined in the previous section, Novae can analyze multiple panels and tissues, with the ability to identify both shared and tissue-specific spatial domains. While we anticipate some overlap in domains across tissues, we also expect certain domains to be specific to particular conditions or diseases. In Figure 2a/c, we present the spatial domains identified across various slides for both human (Figure 2a) and mouse (Figure 2c) tissues. Notably, the lymph node and tonsil exhibited similar spatial domain distributions, with some domains also observed in other tissues such as breast and lung. Also, the heatmap dendrogram regroups slides with similar patterns of domains, and is not a perfect tissue match. Indeed, the spatial domains include also context related to diseases or certain mechanisms, which can be retreived for multiple different tissues. For instance, a lung tumor and a breast tumor may share some cancer-related domains that cannot be found in a healthy breast tissue^20^. The UMAP^21^ representation of spatial context is shown in Figure 2b/d, where each dot corresponds to a cell’s spatial context, clustered into distinct spatial domains. Regarding the mouse study, we considered brain slides, as well as colon, femur, and also a whole mouse sample, as detailed in subsection 4.12. More specifically, the mouse brain slides showed strong inter-slide similarity, as many domains were consistently identified across all slides (see Figure 2c). Additionally, we observed that many domains, including “brain-specific” and “colon-specific” domains, were retrieved in the whole mouse sample. Examples of spatial domains for the whole mouse sample are shown in Figure 2e. For instance, we find the areas characterizing the bones and costal cartilage (D997), the lung lobes (D983), the liver (D973), and the intestine (D975) in the abdominal cavity^22^, as highlighted in Figure 2f. In summary, while expecting results were shown in Figure 2, it shows the capability of Novae to train across multiple tissues, while avoiding over-correction (i.e., the identification of all domains in all slides) or under-correction (i.e., all spatial domains being specific to one slide).

**Figure 2:**
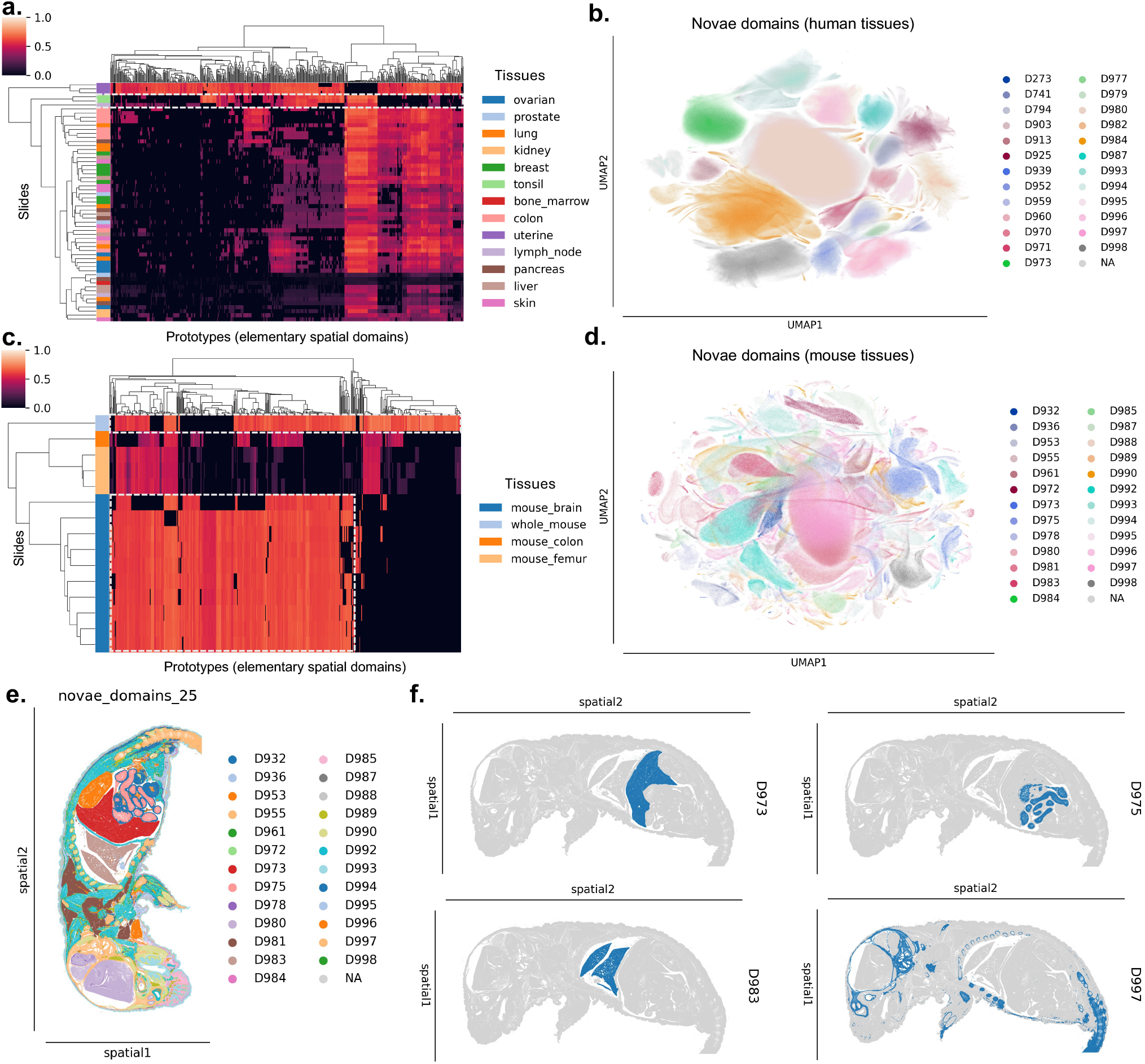
Spatial domains across tissues and species. **a**. Heatmap showing the spatial domain weights for each slide, organized by tissue type. A weight of 0 (black) indicates the absence of the domain in that slide, while higher weights represent increased confidence in the presence of the domain. **b**. UMAP visualization of spatial domain representations in human tissue slides. Each dot corresponds to the spatial context representation of a cell. The panels **c**. and **d**. show similar figures to (a) and (b), but for the mouse samples. **e**. Spatial domains identified for the whole mouse sample. **f**. Zoom-in on four domains in the whole mouse: liver (D973), intestine (D975) within the abdominal cavity, lung lobes (D983), bones and costal cartilage (D997).

### 2.3 High integration and continuity of the spatial domains

In this study, we compared Novae in both zero-shot and fine-tuning modes to four state-of-the-art methods: SpaceFlow^10^, GraphST^9^, SEDR^11^, and STAGATE^8^. We evaluated the performance of these methods across three distinct test cases. In the first test case, we used the breast dataset, composed of two slides with different gene panels. As mentioned earlier, Novae can be trained across multiple gene panels; hence, a single Novae model was trained for both panels. In contrast, the other methods were trained on the intersection of 185 common genes across the two panels (as illustrated in Figure 3a), followed by batch effect correction using Harmony^12^ and clustering with mclust^14^ (using 7, 10, and 15 clusters). To evaluate model performance, we used the FIDE score and Jensen-Shannon Divergence (JSD) score to assess spatial domain continuity and cross-slide homogeneity, respectively (see subsection 4.11 for more details). Figure 3b presents the results of this benchmark, highlighting a significant improvement in performance by Novae, even in the zero-shot case (i.e., using a pre-trained model directly).

**Figure 3:**
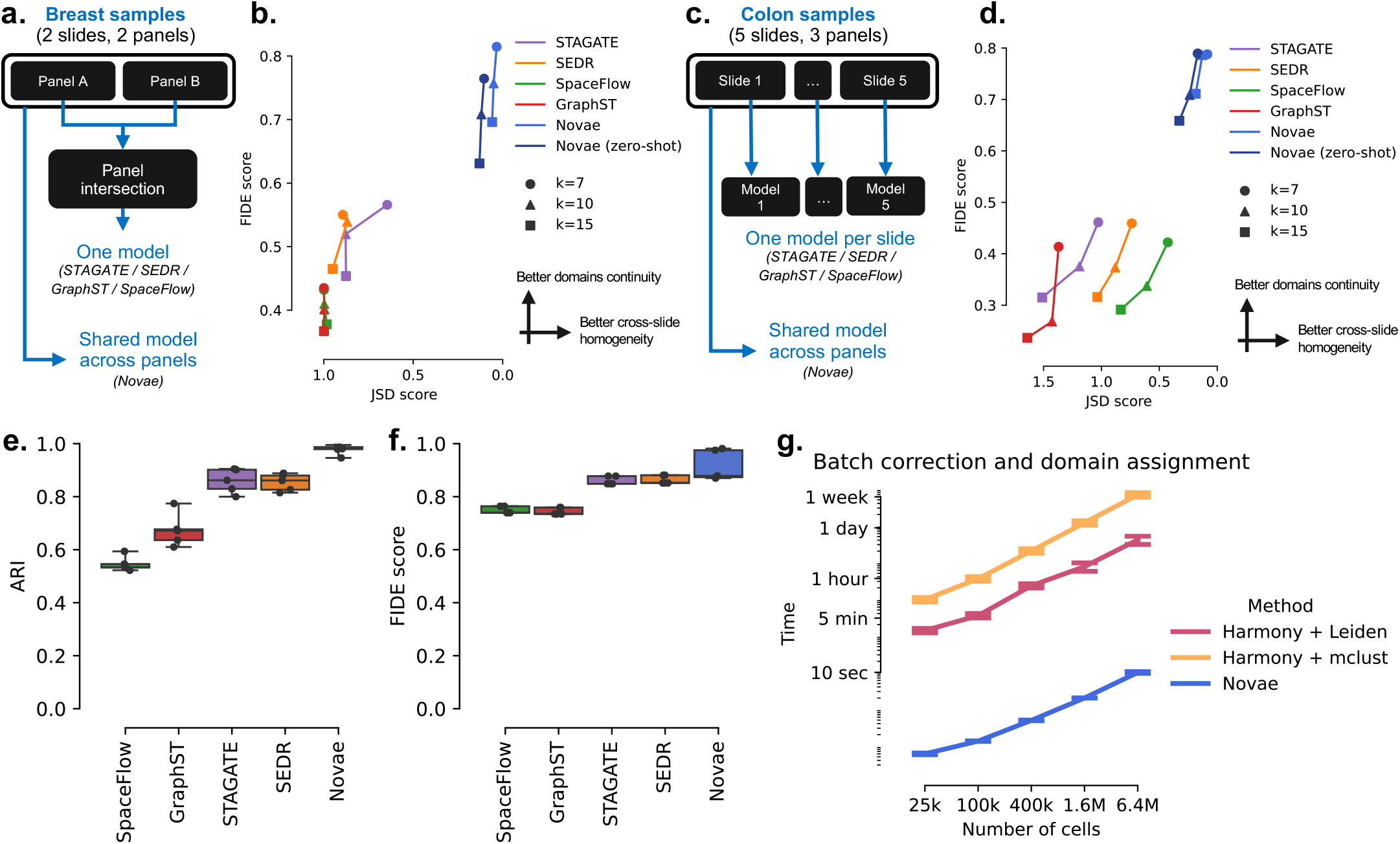
Quantitative benchmark of the spatial domain assignment. **a**. Schematic representation of the benchmarking approach for the breast dataset. The Novae model can be trained on both panels, while other models were run using genes shared between the panels. **b**. FIDE and JSD scores of different methods hree domain counts (7, 10, and 15) on breast samples. The FIDE score evaluates domain continuity, while the JSD score assesses cross-slide homogeneity.**c**. shematic representation of the benchmarking approach for the colon dataset. The same Novae model can be trained on all panels, whereas other models were run a one-per-slide manner. **d**. FIDE and JSD scores of different methods for three domain counts (7, 10, and 15) on colon samples. **e**. ARI comparison on the synthetic dataset. **f**. FIDE score comparison on the synthetic dataset. **g**. Runtime comparison of post-inference tasks (batch-effect correction and domain assignnment) across different dataset sizes.

The second test case involved the colon dataset, where Novae was again trained across all slides. For the other methods, due to the limited intersection of genes between panels, we employed a different approach: a separate model was trained for each slide (as illustrated in Figure 3c). The results from each model were then concatenated, followed by batch-effect correction and clustering. Figure 3d presents the result metrics for all methods on the colon dataset, again showing a superior performance by Novae in both zero-shot and fine-tuning modes.

For visual comparison, Figure 4 shows the spatial domain assignments for the breast samples (see supplementary materials for colon samples). Notably, Figure 4 shows that Novae (i) better identifies cross-slide domains (see the third figure column) and (ii) better aligns the batches/slides in the latent space (see the fourth figure column). Regarding the biological interpretation of the results from Novae, domains D498 and D499 were identified as stromal regions characterized by the expression of *FN1* and *COL1A1* (see supplementals), which are present in both samples. Domain D504 was composed of glandular cells, marked by the expression of *TAPBP, PGR*, and *CDH1*. In the Xenium sample, an expansion of domain D503 was observed, potentially resulting from clonal expansion of cancer cells. This expanded D503 became embedded within the stromal areas of D498 and D499, while excluding the immune cell-rich region D485, which is known for high levels of *PTPRC, CD52*, and *CD3E*. This exclusion pattern, as depicted in the PAGA graphs in the supplementals, is characteristic of tumors with an immune-excluded phenotype.

**Figure 4:**
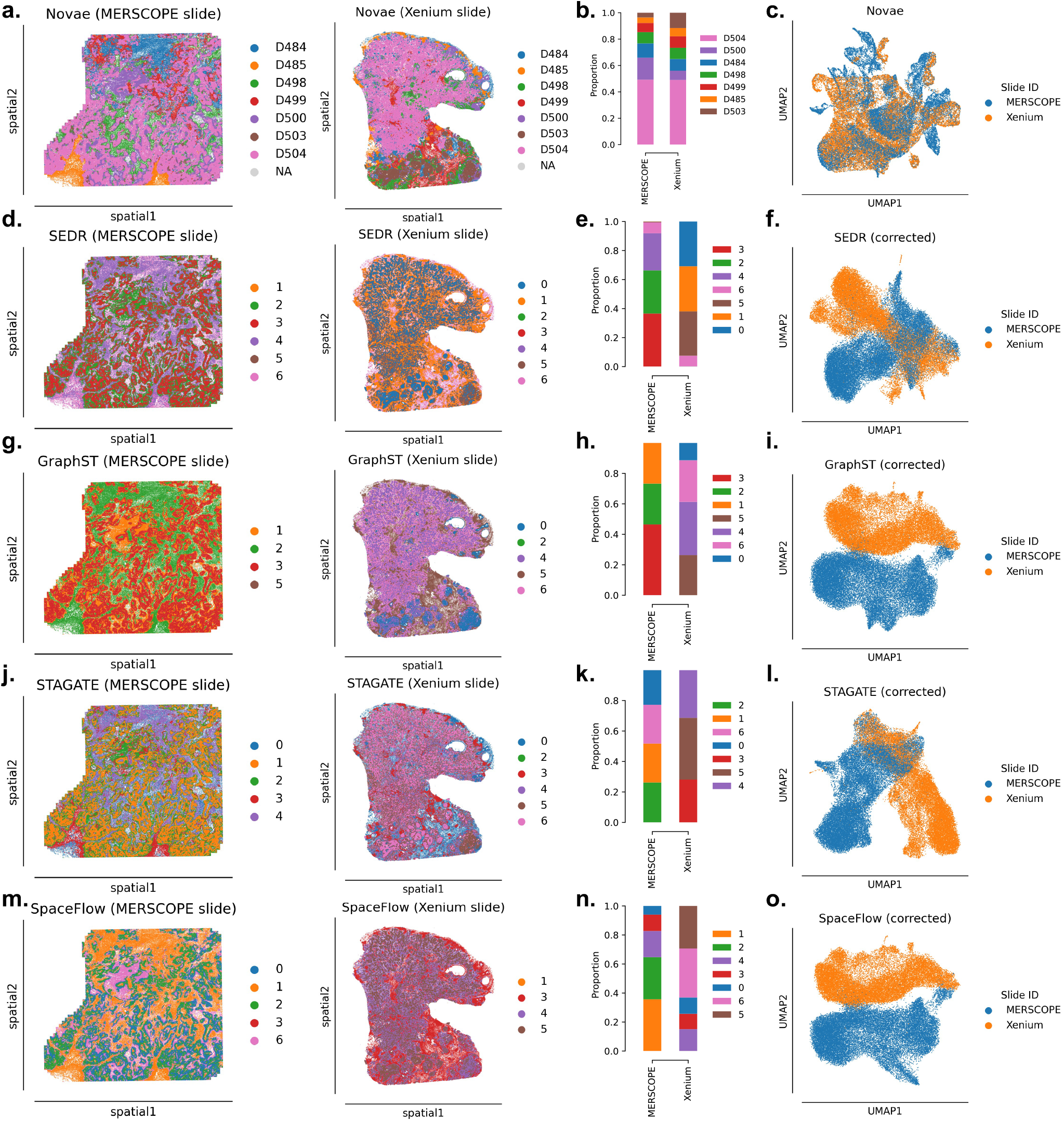
Visualization of the spatial domains for the breast dataset. Spatial domains results across two slides from the breast dataset (Mer-scope and Xenium slides, see more details in subsection 4.12) using five different methods: Novae, SEDR, GraphST, STAGATE, and SpaceFlow. **a**. Spatial domains assigned by Novae on the two breast slides. **b**. Proportions of each Novae domain for each slide. **c**. UMAP of spatial representations, colored by slide ID. Each dot is the UMAP spatial representation of one cell. The next four rows, that is **d**. to **o**., show the same figures as the first row but for the four other methods (SEDR, GraphST, STAGATE, and SpaceFlow, respectively).

The third comparison was conducted on a synthetic dataset consisting of 5 slides and 7 spatial domains (see subsection 4.12 for more details). Unlike the previous cases, this dataset used the same gene panel across all slides, allowing the other methods to run without requiring gene panel intersection. In this case, we did not use Novae in zero-shot mode, as the gene expression was synthetic, meaning the generated cell types may not correspond to real data. The models were evaluated using the Adjusted Rand Index (ARI) for clustering accuracy against the known ground truth, as well as the FIDE score. Figure 3e shows the ARI of the different methods across 5 seeds, while Figure 3f presents the FIDE score across 5 seeds. Again, Novae outperformed the other methods, demonstrating higher ARI and FIDE scores with notably low standard deviation in ARI (Figure 3e).

### 2.4 Time and memory efficiency

After running inference, i.e., computing the cell’s spatial representations, Novae can perform the attribution of the niches and correct batch effect in a short time. Indeed, the attribution of spatial domains is a mapping between the prototypes and the desired resolution (see subsection 4.7 for more details), which is performed in a low (constant) time. Regarding batch-effect correction, since the categorical domain assignments are already corrected through Novae, we can use the assignments to align the spatial representations (as detailed in subsection 4.9). This vectorial operation is also fast and in linear time. Contrastively, for the two latter operations, the other state-of-the-art models depend on external tools: usually (i) Harmony^12^ for batch-effect correction and (ii) Leiden^13^ or mclust^14^ to assign spatial domains to the representations. As shown by Figure 3g, this can be slow, especially on large datasets of millions of cells. Indeed, this can take up to several days on 6 millions cells, while Novae can perform these two operations in several seconds. Furthermore, during experimentation, it is common to try multiple resolutions of spatial domains, hence requiring clustering to be run multiple times. Novae can perform this very rapidly, thus easing the analysis of different resolutions, while the other methods require running a time-consuming clustering again. Also, regarding random-access-memory (RAM) usage, Novae supports lazy loading. That is, instead of storing the full graph dataset in memory, each subgraph is created on the fly before running through the model. This prevents a significant amount of RAM from being dedicated to loading the dataset. This allowed us to train Novae on a dataset composed of nearly 30 million cells using a GPU with 40GB of RAM (see subsection 4.14 for more details). In addition, Novae runs on local subgraphs instead of running on a full slide, which is original for two reasons. First, the subgraphs are smaller and therefore require less memory. Secondly, it is possible to use a larger number of layers in the neural network without aggregating information from further environments. For instance, having a graph neural network of 16 layers operating on the full slide would lead to mixed information between two cells at a distance of 300 microns, which is usually not intended. Instead, using local subgraphs ensures only aggregating information from local microenvironments.

### 2.5 A multitude of downstream tasks

Novae’s spatial representations enable the performance of multiple downstream tasks. To illustrate this, we applied three tasks across two datasets. The first dataset consists of a non-diseased lymph node and a reactive lymph node (shown in panels Figure 5a-d), while the second dataset comprises six mouse brain slides collected at different time points, with half exhibiting Alzheimer’s-like pathology (further details in subsection 4.12). For the lymph node dataset, spatial domains are displayed in Figure 5a, with Figure 5b/c providing details of gene expression and domain proportions to help the characterisation of these spatial domains. To compare the spatial organization (or “slide architecture”), users can apply trajectory inference methods to the spatial representations generated by Novae (more details in subsection 4.10). As shown in Figure 5d, we observed changes in the spatial organization of these domains. Notably, the D500 domain in the germinal center, enriched with (*CD79*+ *CR2*+) mature B cells, was previously connected only to the D501 domain before clonal expansion. Additionally, Figure 5c reveals an inversion in the proportions of the D500 and D501 domains, also visible in Figure 5a.

**Figure 5:**
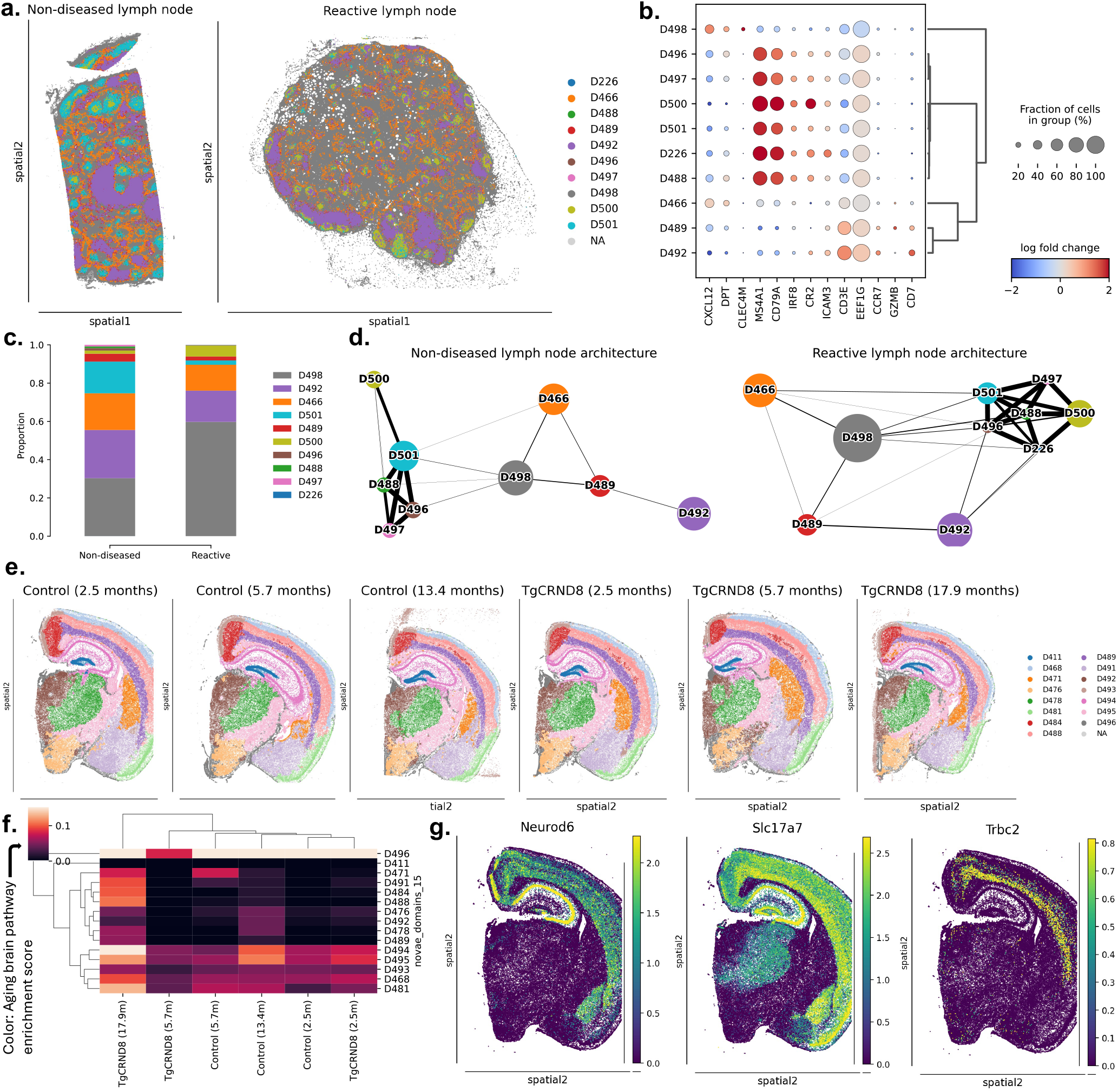
Downstream tasks examples on human lymph node and mouse brain slides. **a**. Spatial domains of the non-diseased (left) and reactive (right) lymph nodes. **b**. DEGs between the spatial domains of the reactive lymph node slide. **c**. Proportions of spatial domains for each lymph node slide. **d**. PAGA graphs comparing the domain organization of non-diseased (left) and reactive (right) lymph nodes. **e**. Spatial domains of mouse brain across different time points, comparing control conditions and Alzheimer-like pathology (*TgCRND8* samples). **f**. Heatmap of brain aging pathway activation across different domains and time points in the mouse brain slides. **f**. Expression patterns of three spatially variable genes on the 2.5-month control mouse brain slide.

Regarding the mouse brain dataset, the spatial domains identified by Novae are presented in Figure 5e. Although no significant changes in brain architecture were observed (see supplementals for details), we do observe differences in brain aging pathway^23^ enrichment that we computed for each spatial domain. As shown in Figure 5f, the 17.9-month-old *TgCRND8* mouse, which exhibits Alzheimer’s-like pathology, shows higher brain aging, particularly in specific spatial domains such as D494 and D481. For example, the D494 domain has a high expression of the *Neurod6* gene, a key gene associated with brain aging^23^. This analysis highlights how identifying spatial domains enhances the understanding of pathway activation, demonstrating the utility of spatial domain identification in pathway analysis. In addition, differential gene expression analysis can be performed on cells grouped by spatial domains, allowing for the identification of spatially variable genes (SVGs). SVGs are genes whose expression varies significantly across spatial domains. In Figure 5g, we demonstrate this for the 2.5-month control mouse, showing the three most spatially variable genes identified by Novae. Therefore, this helps better identify genes that have different activations depending on the spatial domain.

## 3 Discussion

The field of spatial transcriptomics is advancing rapidly, leading to the generation of increasingly larger datasets^24^. While this growth holds the promise of uncovering new insights that were difficult to achieve with scRNAseq^25^, it also presents significant challenges in performing comparative spatial analyses across multiple slides. Current algorithms^8–11^ are often designed for single or consecutive slide analyses and typically rely on external batch-effect correction methods, which may not be sufficient for the complexities of multi-slide spatial datasets. Novae addresses this gap by offering a more flexible solution that can operate across various tissues, conditions, technologies, and gene panels. Compared to related models, we have shown that Novae can better identify cross-slide domains while not over-correcting the spatial domain assignments. Indeed, the flexibility introduced in the prototypes allows for certain domains to be tissue-specific or disease-specific. In addition, Novae is less dependent on external tools, as it performs domain assignment and batch-effect correction inside the same model. This has many advantages, such as (i) a better batch-effect correction due to the spatial awareness of the model and (ii) improved speed performances. Also, as a shared open-source model, Novae can be easily loaded and used for user-specific tasks without the need for hyperparameter optimization or large re-training.

Our study demonstrates Novae’s potential, particularly when applied to large cohorts, to identify spatial domains enriched or specific to certain conditions or diseases. Using spatial trajectory analysis, we demonstrated how tissue organization and structure can be characterized, as shown by the comparison of a reactive lymph node with a non-diseased one. Lymph nodes are crucial for immune surveillance across different organs, with their structure enabling a targeted immune response during infections. In our analysis, we observed spatial reorganization in the reactive lymph node, notably the expansion of the D498 domain, which is enriched in the chemoattractant CXCL12, a key factor in recruiting inflammatory cells. Concerning the second use case example, we identified an enriched aging-related pathway in specific mouse brain regions linked to Alzheimer’s-like pathology at defined time points, as well as brain-specific spatially variable genes. Overall, this study has showcased Novae’s versatility in handling multiple downstream tasks, such as spatial domain identification, slide architecture, and spatially variable gene analysis, all within a single framework.

Methodologically, Novae is built upon the SwAV^18^ framework, a self-supervision technique initially developed for computer vision. In this study, SwAV was adapted for graph learning, and many components were updated to better suit our biological problem, e.g., using biologically relevant augmentations. Furthermore, compared to other self-supervised methods, SwAV has two major properties that make it suitable for our purpose. First, SwAV’s inclusion of optimal transport^19^ (OT) within the model is particularly noteworthy, as OT has proven effective in addressing batch effects natively, offering a significant advantage over traditional methods that require external corrections. Secondly, SwAV’s approach to learning embeddings in the latent space (that is, prototypes) allows for efficient assignment of spatial domains, with prototypes serving as centroids for elementary classes. Since we typically have less than a thousand prototypes, hierarchical clustering on prototypes is computationally efficient and facilitates the definition of nested spatial domains, providing an advantage over more conventional clustering techniques on the latent space.

Integrating additional modalities into Novae, for instance, protein data, could lead to the development of a multi-omics spatial model. This could be achieved by incorporating protein embeddings into the model, allowing it to analyze both transcriptomic and proteomic data concurrently. Another area for improvement lies in the segmentation process of our training dataset, which currently relies on proprietary methods that may benefit from refinement. Given the critical role of segmentation in determining the quality of spatial data^26^, the application of newer segmentation tools to the raw data could enhance overall dataset quality. Additionally, using multiple segmentation methods could serve as a form of data augmentation, potentially improving the model’s robustness. Addressing these areas will further enhance Novae’s utility and effectiveness in spatial transcriptomics research.

## 4 Online Methods

### 4.1 Model input

In Novae, we consider a total of *G* genes, such as those in the human or mouse genome. A spatial transcriptomics slide captures the expression profile of a subset of these genes, denoted by 𝒫 ⊆ {1, …, *G*}, representing the indices of the genes in this specific panel. Throughout this article, we use the term “cell” to refer to either a spot or a cell, although Novae is applicable to both spot-resolution and single-cell resolution technologies. The expression data for a single slide is represented by a matrix **X** = (**x**_1_, …, **x**_*N*_) ∈ ℝ^*N*×*P*^, where *N* denotes the number of cells and *P* = Card(𝒫) represents the number of genes in the panel P. This matrix **X** contains normalized and logarithmized gene expression values. To incorporate spatial information, we utilize the 2D localization of cells to construct a Delaunay graph. In this graph, nodes correspond to cells, and edges indicate neighboring cells. The adjacency matrix **A** ∈ ℝ^*N*×*N*^ encodes the connectivity between cells, with edge weights representing the distances between neighboring cells in the Delaunay graph. Also, an entire spatial transcriptomics dataset consists of multiple slides, each described by a tuple (**X**, 𝒫, **A**), where the size of each gene panel Card(𝒫) can vary across different slides. Additionally, for a given slide and a specific cell *i*, we define a subgraph G_*i*_ consisting of cells within a distance of *n*_*local*_ edges from cell *i*. The features of a node (or cell) in this subgraph are the gene expression vectors of the corresponding cells. For instance, the feature vector for cell *i* is **x**_*i*_ ∈ ℝ^*P*^. In the following sections, 𝒢_*i*_ is used to obtain a spatial domain representation for the subgraph origin cell *i*.

### 4.2 Augmentation

Augmentation is a technique used to introduce artificial variations into the data, enhancing the dataset’s diversity. This approach is commonly employed to improve the robustness and generalization abilities of models. In the context of spatial transcriptomics, we apply two biologically relevant augmentations to strengthen the robustness of Novae.

Firstly, we introduce a “pseudo batch effect” noise to reduce the model’s sensitivity to batch effects. For a given subgraph input, we sample two vectors: **a** ∼ exponential(*λ*)^*P*^ and **s** ∼ Normal(0, *σ*^2^**I**_*P*_), where *λ* and *σ* are hyperparameters. The gene expression vector for each cell *i* is then updated as 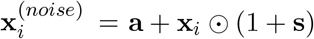.

Secondly, we randomly subset the gene panel according to a ratio *γ* ∈]0, 1[. For a given panel 𝒫, we select ⌊*γP*⌋ genes, resulting in a new panel 𝒫′ ⊂ 𝒫. This augmentation simulates the effect of different panels of genes, as if multiple machines generated the data, or as if the panel was updated during a study. After applying these augmentations, each cell is represented by the expression vector 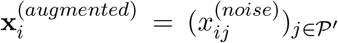. This augmented representation helps the model better generalize across varying conditions and datasets.

### 4.3 Cell embedding

After augmentation, cells are transformed into panel-invariant embeddings, which serve as the node features for the graph encoder described in the subsequent section. Let **v**_**1**_, …, **v**_**G**_ ∈ ℝ^*E*^ denote a list of trainable gene embeddings, where *E* represents the embedding size. For a given gene panel 𝒫 ⊆ {1, …, *G*}, we consider only the embeddings corresponding to the genes in this panel. These embeddings are L2-normalized and then multiplied by the cell’s gene expression vector **x** ∈ ℝ^*P*^. Specifically, the embedding for a cell *i* is calculated as follows:

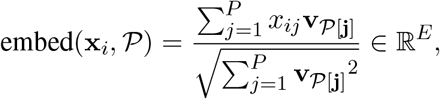

where the square root, square, and division operations are performed element-wise. The purpose of the L2 normalization is to ensure that the embedding weights are comparable across different gene panel sizes. This can be compared to a principal component analysis (PCA) reduction, where the components are trainable gene embeddings, and the L2 normalization ensures that each panel has consistent weights across different “gene programs”. Specifically, for each *e* ≤ *E*, **v**_*ge*_ represents the weight of gene **v**_*g*_ in the gene program *e*. Additionally, rather than training all gene embeddings from scratch, we can initialize with pre-trained gene embeddings from scGPT^27^. To prevent domain shift for genes not present in the Novae dataset, these pre-trained embeddings are frozen, and we introduce a trainable linear layer afterward. Thus, the updated cell embedding formula becomes

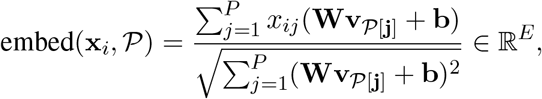

where **W** ∈ ℝ^*E*×*E*^ and **b** ∈ ℝ^*E*^ are trainable parameters. During training, the function embed(·, ·) is applied to the augmented gene expression vector 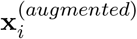, whereas during prediction or inference, it is applied to the original gene expression vector **x**_*i*_.

### 4.4 Graph encoder

The graph encoder utilized in Novae is a Graph Attention Network^16^ (GAT), which employs attention mechanisms to aggregate information from neighboring cells. The GAT is composed of multiple layers, with each layer potentially having multiple attention heads. The input to the GAT is a subgraph G, where node features are embedded cell features, as described in subsection 4.3. For each cell *i*, the initial node feature is 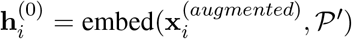 during training, and 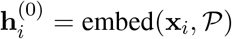 during inference. For each layer *l*, the node features for the next layer are calculated as

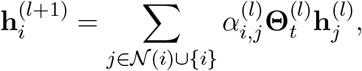

where the attention coefficients 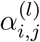 are defined as

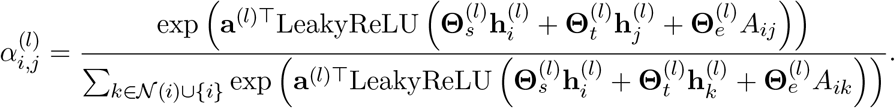

In these equations, 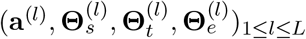 denotes parameters of the model (biases or matrices), and 𝒩 represents the set of neighbors of a cell in the subgraph 𝒢. After *L* sequential layers, each node *i* has a feature vector 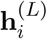. Then, to obtain a unified representation for the entire subgraph 𝒢, we employ an attention aggregation layer, resulting in a graph-level representation **z** := attention-aggregation 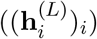. The subgraph representation **z** is learned through the self-supervised task described in subsection 4.5. Note that, for simplification, the above equations detail the graph encoder with only one head. With multiple attention heads, the GAT computes separate attention coefficients and node features for each head. The outputs from all heads are then concatenated (or averaged) to form the final node representation for each layer, allowing the model to capture diverse aspects of the node’s neighborhood.

### 4.5 Prototypes and swapped assignment task

Since the dataset lacks ground truth, we need to train Novae using an unsupervised approach. Specifically, as we aim to pre-train a foundation model, we will leverage self-supervised learning^15,28^, which is well-suited for capturing meaningful data representations. Among the different self-supervision frameworks, SwAV^18^ is a self-supervised learning algorithm that integrates contrastive learning and clustering. The main concept is to learn representations by predicting cluster assignments derived from different augmentations (views) of the same image. In our context, instead of using two views of the same image, we utilize two closely related subgraphs of cells. Specifically, we select a pair (*i, j*) of cells separated by *n*_*view*_ edges, from which we derive the corresponding subgraphs (𝒢_*i*_, 𝒢_*j*_). When *n*_*view*_ is small enough (typically 2 or 3), we expect that the subgraph representations (**z**_*i*_, **z**_*j*_) will belong to the same spatial domain. To actually assign these representations to spatial domains, we train a set of *K* vectors **c**_1_, …, **c**_*K*_ ∈ ℝ^*O*^ in the unit sphere (i.e., 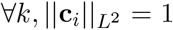, where 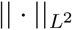 is the L2-norm). These trainable embeddings are referred to as *prototypes* in the original SwAV^18^ paper. In our application, they represent high-resolution spatial domain (i.e., in practice, *K* is large). The representation **z**_*i*_ of each cell is projected onto the prototypes, over which a softmax function is applied:

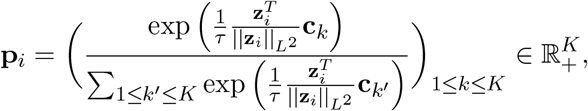

where *τ* is a temperature parameter that controls the sharpness of the softmax distribution. Intuitively, *p*_*ik*_ represents the probability that cell *i* belongs to prototype *k*. To prevent the representations from collapsing into a single prototype, we define a “corrected” assignment **q**_*i*_ that aligns with the distribution **p**_*i*_ while considering global mini-batch statistics. Essentially, we aim for each mini-batch to represent all prototypes as evenly as possible. This assignment **q**_*i*_ is derived from the result of an optimal transport (OT) problem over a mini-batch of size *B* (with each mini-batch being dedicated to one slide). Specifically, given a mini-batch of *B* representations **Z** = (**z**_1_, …, **z**_*B*_) ∈ ℝ^*B*×*O*^ and the matrix of all prototypes **C** = (**c**_1_, …, **c**_*K*_) ∈ ℝ^*K*×*O*^, the OT problem is defined as:

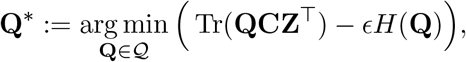

where *H* is the Shannon entropy, *ϵ* is a regularization hyperparameter, and 𝒬 is the transportation polytope defined by 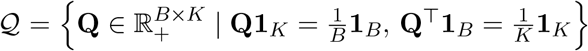. This OT problem ensures a smooth distribution of assignments across the different prototypes, preventing mode collapse. It also helps avoid learning “artificial” prototypes that are specific to certain slides or batch effects. The Sinkhorn-Knopp algorithm is employed to approximate **Q**^*^, as detailed in the supplementary notes. Subsequently, we obtain 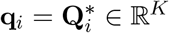, which represents the “corrected” assignments for cell *i*. The loss function used in Novae is a cross-entropy applied to the “swapped” prediction problem. Specifically, **q**_*i*_ serves as the “self-supervised ground truth” for **p**_*j*_, while **q**_*j*_ is used for **p**_*i*_. We formally define the loss as:

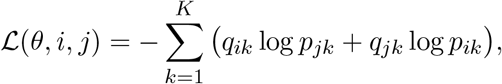

where *θ* represents all the model parameters, and (*i, j*) denotes a pair of cells separated by *n*_*view*_ edges. This loss function encourages the soft spatial domains of cell *i* to resemble the soft “pseudo-ground-truth” spatial domains of cell *j*, and vice versa. The model is trained via backpropagation of this loss using the Adam optimizer. Note that the gradients are detached during the optimal transport part, meaning that the loss is backpropagating via (**p**_*i*_, **p**_*j*_).

### 4.6 Pan-tissue prototypes

The loss defined in the previous section is limited when training on multiple tissues, as the optimal transport problem enforces similar distribution assignments to the prototypes across all slides. However, many tissues possess unique, non-overlapping spatial domains, making it biologically unrealistic to train shared prototypes for all tissues. To address this, we introduce a slide-by-prototype matrix of weights, allowing the model to not map a slide to all domains. Specifically, we initialize a queue **W**^(*queue*)^ ∈ ℝ^*S*×*size*×*K*^, where *S* is the number of different slides, *size* is the queue size, and *K* is the number of prototypes (filled with 1*/K* everywhere). For each slide and mini-batch of size *B* (the mini-batch contains cells of the same slide), we calculate the maximum assignment probability for each prototype, 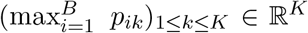, and store this in the queue at the corresponding slide’s position. Technically, this involves rolling the queue weights for a specific slide *s* and updating the entry as 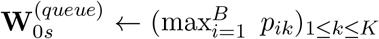. Afterward, we compute the maximum probabilities over the queue size, i.e. the second dimension, which we denote **W**^*^ ∈ ℝ^*S*×*K*^. This slide-by-prototype matrix **W**^*^ is used for each mini-batch to subset the prototypes on which the optimal transport is performed to define the corrected assignments **q**. For a given slide *s* ∈ {1, …, *S*}, the codes are computed using the prototypes *k* for which 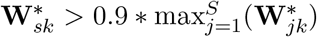, meaning that the probability of having this prototype is high enough (compared to the slide which most likely have this domain). Yet, in order to not only learn slide-specific prototypes, we ensure that each slide is mapped over at least *ρK* prototypes, where *ρ* ∈]0, 1[ is a hyperparameter. If less than *ρK* prototypes match the above condition, we choose the ⌊*ρK*⌋ prototypes with the highest 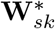 score. Typically, we choose 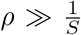 to ensure that prototypes will be shared by multiple slides. Indeed, we expect slides from the same tissue to have shared prototypes (or “tissue-specific” prototypes), while some prototypes should be shared across multiple tissues.

### 4.7 Assignment to spatial domains

Prototypes can be considered as centroids of elementary spatial domains. Since the desired number of spatial domains may vary, we employ hierarchical clustering on the prototypes. In this setup, the prototypes serve as the leaves of a hierarchical tree, with each level of the tree representing increasingly coarse-grained spatial domains. For a given subgraph representation **z**_*i*_, we assign it to the closest prototype by selecting the one with the highest dot product, defined as 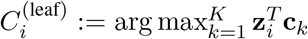. This assignment 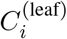 indicates to which tree-leaf the cell *i* is associated. At a particular level *l* of the tree, the spatial domain assignment for cell *i* is determined by 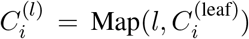, where Map(*l*, ·) is a mapping function that associates the leaf prototype 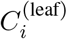 with a cluster at level *l* (the level *l* is chosen according to the number of desired clusters). This hierarchical approach allows each cell representation to be assigned to spatial domains of varying resolutions efficiently (constant time). In other words, the prototypes serve as elementary spatial domains, used to define spatial domains at the desired resolution.

### 4.8 Zero-shot and fine-tuning

We train Novae on a large dataset composed of 18 different tissues, as detailed in subsection 4.12. Therefore, we can save this model on a hub (in particular, Hugging Face Hub), and then anyone can download this model and re-use it on their own dataset without re-training. For that, for a given new dataset, we compute the spatial domain representation **z** of all cells in all slides. Afterward, we run a KMeans algorithm from sklearn^29^ on the representations to define *K* centroids, which will act as the new prototypes. These proto-types are used for spatial domain assignment as detailed in the previous section. We denote this as zero-shot since Novae is not re-trained. For fine-tuning, we apply the same approach but then re-train the model for a few epochs, potentially one only epoch.

### 4.9 Batch effect correction

The optimal transport approach ensures the retrieval of relatively similar spatial domains across slides of the same tissue. Therefore, the spatial domain assignments 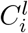 are corrected for the different slides, meaning that correcting the actual representations **z** is not always necessary. If correcting the representation is actually desired, for each spatial domain we compute the centroid representation of the slide that has the most cells in this spatial domain, which generates a list of centroids 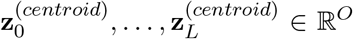. Then, for each slide and for each spatial domain *l*, we compute the mean representation 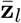, and the representation of a cell *I* in spatial domain *l* becomes 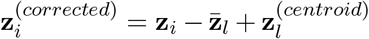. Essentially, we translate the representations to align the centroids to the reference centroid.

### 4.10 Downstream tasks examples

For downstream tasks, we either use the spatial representations of Novae (i.e., vectors), or directly the spatial domain assignments per cells (i.e., categories). For instance, the spatially variable genes (SVG) are defined by the `scanpy.tl.rank genes groups` function from Scanpy^30^ over the categorical spatial domains of Novae, and increasing the resolution of the Novae domains results in more local SVG. Regarding pathways, we use the `scanpy.tl.score genes` function from Scanpy applied to gene sets from the Gene Set Enrichment Analysis (GSEA) database^31,32^. This gives a score for every cell and every pathway. Afterward, we average these scores per spatial domain, resulting in a domain-by-pathway heatmap. We plot this heatmap using the `seaborn.clustermap` function from Seaborn^33^, which groups domains with similar patterns of scores across pathways. More specifically, in Figure 5, we used the LEE AGING CEREBELLUM UP^23^ pathway (https://www.gsea-msigdb.org/gsea/msigdb/mouse/geneset/LEE_AGING_CEREBELLUM_UP.html?ex=1). Regarding trajectory inference, we can run any method on the representations of Novae, grouped by categorical spatial domain. In practice, in this article, we used PAGA^34^ and more specifically the `scanpy.tl.paga` implementation from Scanpy.

### 4.11 Metrics for model comparison

The F1-score of inter-domain edges (FIDE score) is used to quantify the spatial domain continuity. More specifically, let **C** = (*C*_*i*_)_1≤*i*≤*N*_ be the categorical spatial domain predictions for *N* cells of a slide. The FIDE score is defined as FIDE(**C, A**) = F1-score((*C*_*i*_, *C*_*j*_)_*i,j* s.t._*A* _*ij*_>0), where A_*ij*_ is positive when the cells *i* and *j* are graph neighbors. Intuitively, for an edge *i* ↔ *j*, the edge is an inter-domain edge if *C*_*i*_ ≠ *C*_*j*_. A high number of inter-domain edges shows a low domain continuity, therefore quantified by a low FIDE score. On multiple slides, we average the per-slide FIDE scores. The Jensen-Shannon divergence (JSD) is used to quantify the homogeneity of domains across multiple slides. For each slide *s* ∈ ⟬1, *S* ⟭, we compute *π*_*s*_ ∈ [0, 1]^*D*^, the proportion of each domain among the slide, where *D* is the total number of spatial domains. Then, the JSD metric is defined as 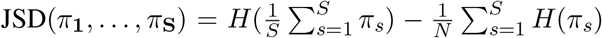, where *H* is the Shannon entropy. The Adjusted Rand Index^35^ (ARI) is defined as the percentage of concordance between two clusterings. Mathematically, 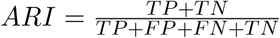, where *TP* is the number of true positives, *TN* the number of true negatives, *FP* the number of false positives, and *FN* the number of false negatives.

### 4.12 Datasets used

In total, 78 public spatial transcriptomics slides were collected from single-cell resolution technologies (Xenium, MERSCOPE, CosMX). The datasets are available on the vendor websites, i.e., respectively 10X Genomics, Vizgen and Nanostring. This dataset covers 19 tissues (human bone marrow, human brain, human breast, human colon, human kidney, human liver, human lung, human lymph node, human ovarian, human pancreas, human prostate, human skin, human tonsil, human uterine, mouse femur, mouse brain, mouse colon, and whole mouse) and a total of 28,909,516 cells. It is comprised of 853,622 cells from the CosMX technology, 10,196,835 cells from the MERSCOPE technology, and 17,859,059 cells from the Xenium technology. The CosMX covers 2 different tissues, 8 for the MERSCOPE, and 17 for the Xenium technology. In total, 8 tissues contain slides from multiple technologies (breast, colon, liver, lung, ovarian, pancreas, prostate, and skin). For the benchmark, we selected five colon slides from the Xenium technology, divided into three distinct gene panels. The first panel includes 325 genes and comprises 270,984 cells, the second panel contains 422 genes and 924,597 cells, and the third panel consists of 480 genes, with a total of 388,175 cells. The slides can be found in the Novae dataset (see data availability statement), and their name is listed in Supplementary Table 1. We also selected two breast slides from different platforms: Xenium and Merscope, containing a total of 1,290,084 cells. Again, their name is listed in Supplementary Table 1. Similarly, we listed in Supplementary Table 1 the name of the lymph node and mouse brain slides used in Figure 5. Regarding the mouse brain slides, note that the “Alzheimer-like pathology” concerns the CRND8 APP-overexpressing (TgCRND8) transgenic mouses, as detailed in https://www.10xgenomics.com/datasets/xenium-in-situ-analysis-of-alzheimers-disease-mouse-model-brain-coronal-sections-from-one-hemisphere-over-a-time-course-1-standard. Additionally, we created a synthetic dataset in order to compare the different models to a ground-truth. More specifically, 7 domains were generated in a circular manner, for 5 different slides with 100 genes. For each domain of each slide, the gene expression was sampled from an exponential law whose parameter is the sum of a slide-specific parameter and a domain-specific marker. For more details, see the supplementals.

### 4.13 Spatial-domains assignment benchmark

In our benchmark study, we evaluated the performance of our method, Novae, against four state-of-the-art methods: SpaceFlow^10^, GraphST^9^, STAGATE^8^, and SEDR^11^, using three spatial transcriptomics datasets: the colon dataset, the breast dataset, and a synthetic dataset, as described in subsection 4.12. Regarding the colon dataset, each method was applied independently to the different gene panels (except Novae, which works across different gene panels). For the breast dataset, we focused on the intersection of the two slides, which included 185 common genes, ensuring that all methods were applied consistently to this shared gene set. Again, this excludes Novae, which can work on both gene panels at the same time. After training the different models on each dataset, we applied Harmony^12^ to address potential batch effects introduced by the use of different slides or experimental conditions, aligning the spatial transcriptomic data across slides. Following Harmony, we used the mclust^14^ algorithm for clustering the representation, each cluster being considered as a spatial domain. Note that clustering was performed on the concatenated data to get consistent clusters across slides. Again, the usage of Harmony and mclust does not concern Novae, which includes both operations natively, as described in subsection 4.9 and subsection 4.7.

### 4.14 Implementation and training details

The input of Novae is composed of one or many AnnData^36^ objects from the anndata^36^ data structure of the scverse^37^ community. They contain cell-by-gene tables, which can be typically obtained via (i) reading the proprietary data, e.g., using the readers from SpatialData^38^, or (ii) performing a new segmentation, e.g., using CellPose^39^ or Baysor^40^. When constructing the Delaunay graph, edges longer than 100 microns were dropped, which ensures long-distance cells are not considered neighbors. We implemented Novae using Python and the deep learning framework Pytorch^41^ as well as Pytorch Geometric^42^. Each mini-batch contains *B* subgraphs such that all these subgraphs come from the same slide. For RAM efficiency, the subgraphs are lazy-loaded when sampling a mini-batch. The slide queue **W**^(*queue*)^ is not used during the first epochs to ensure it is filled, and the prototypes are also frozen during these epochs. The model training was monitored with Weight & Biases. The hyperparameters were fined-tuned with the sweep option from Weight & Biases on a heuristic, the product of the FIDE score and the Shannon entropy of the spatial domains distribution. Novae was trained on a Nvidia HGX A100 GPU for 24 hours.

## Acknowledgement

This work is supported by (i) IHU Prism funded by the France 2030 programme and the French National Research Agency (ANR) under grant number ANR-23-IAHU-0002, as well as (ii) the Fondation ARC pour la recherche sur le cancer.

## Code availability

The code developed in this article is available as an open-source Python package, accessible on Github at https://github.com/MICS-Lab/novae, and installable via PyPI. The code used to run the benchmark is available at https://github.com/MICS-Lab/novae_benchmark.

## Data availability

The MERSCOPE datasets are available online at https://vizgen.com/data-release-program/. The Xenium datasets are available online at https://www.10xgenomics.com/datasets. The CosMX datasets are available online at https://nanostring.com/products/cosmx-spatial-molecular-imager/ffpe-dataset/. The scripts to download all the datasets are also available in the repository of Novae at https://github.com/MICS-Lab/novae/tree/main/data.

## Contribution statement

Q.B. defined the objectives of the project, defined the methodological approach, implemented Novae, and wrote the manuscript. H.B. performed the computational benchmark between Novae and the existing state-of-the-art models, gave ideas for the methodological approach, and contributed to writing the manuscript. N.B. validated the biological results of the model, and wrote the section related to these findings. P.H.C supervised Q.B., concentrating on the methodological aspects of the work and manuscript development. Finally, F.A. and P.H.C. acquired the grant to fund the project.

## Notes

https://github.com/MICS-Lab/novae

### Competing Interest Statement

The authors have declared no competing interest.

